# Translational mapping of spatially resolved transcriptomes in human and mouse pulmonary fibrosis

**DOI:** 10.1101/2023.12.21.572330

**Authors:** Lovisa Franzén, Martina Olsson Lindvall, Michael Hühn, Victoria Ptasinski, Laura Setyo, Benjamin Keith, Astrid Collin, Steven Oag, Thomas Volckaert, Annika Borde, Joakim Lundeberg, Julia Lindgren, Graham Belfield, Sonya Jackson, Anna Ollerstam, Marianna Stamou, Patrik L Ståhl, Jorrit J Hornberg

**Affiliations:** Safety Sciences, Clinical Pharmacology & Safety Sciences, R&D, AstraZeneca, Gothenburg, Sweden; Department of Gene Technology, KTH Royal Institute of Technology, Science for Life Laboratory, Stockholm, Sweden; Translational Science and Experimental Medicine, Research and Early Development, Respiratory & Immunology, BioPharmaceuticals R&D, AstraZeneca, Gothenburg, Sweden; Pathology, Clinical Pharmacology & Safety Sciences, R&D, AstraZeneca, Cambridge, UK; Quantitative Biology, Discovery Sciences, R&D, AstraZeneca, Gothenburg, Sweden; Animal Science & Technologies, Clinical Pharmacology & Safety Sciences, R&D, AstraZeneca, Gothenburg, Sweden; Bioscience In Vivo, Research and Early Development, Respiratory & Immunology, BioPharmaceuticals R&D, AstraZeneca, Gothenburg, Sweden; Translational Genomics, Centre for Genomics Research, Discovery Sciences, R&D, AstraZeneca, Gothenburg, Sweden; Late-Stage Development, Respiratory & Immunology, BioPharmaceuticals R&D, AstraZeneca, Gothenburg, Sweden

**Author notes:** Equal contribution (Lovisa Franzén, Martina Olsson Lindvall). Correspondence to: Marianna Stamou,; Patrik L Ståhl.

## Abstract

Idiopathic pulmonary fibrosis (IPF) is a progressive lung disease with poor prognosis and limited treatment options. Efforts to identify effective treatments are thwarted by limited understanding of IPF pathogenesis and poor translatability of available preclinical models. To address these limitations, we generated spatially resolved transcriptome maps of human IPF and bleomycin-induced mouse lung fibrosis. We uncovered distinct fibrotic niches in the IPF lung, characterized by aberrant alveolar epithelial cells in a microenvironment dominated by TGFβ signaling alongside factors such as p53 and ApoE. We also identified a clear divergence between the arrested alveolar regeneration in the IPF fibrotic niches, and the active tissue repair in the acutely fibrotic mouse lung. Our study offers in-depth insights into the IPF transcriptional landscape and proposes alveolar regeneration as a promising therapeutic strategy for IPF.

---

Idiopathic pulmonary fibrosis (IPF) is a chronic lung disease characterized by progressive and irreversible scarring of the lung. Treatment options are limited, and the development of new therapies is impeded by incomplete understanding of disease pathogenesis and translatability limitations of available pre-clinical models. Recent advances into mechanistic understanding of IPF pathogenesis reveal complex gene-environment interactions as key pathophysiological drivers^1–3^.

Single-cell studies have revealed IPF-associated cell states, including atypical epithelial cells, fibroblasts^4,5^, and pro-fibrotic alveolar macrophages^6,7^. Interestingly, a novel KRT5-/KRT17+ aberrant basaloid (AbBa) epithelial cell population has been independently identified in multiple studies^4,5,8,9^, expressing epithelial, basal, and mesenchymal markers and genes related to senescence and extracellular matrix (ECM) production. These cells likely originate from alveolar type 2 (AT2) or club cells^4,5,9,10^, but their role in the fibrotic microenvironment remains elusive. A closely related Krt8+ alveolar differentiation intermediate (ADI) cell population is present in the widely used mouse model of bleomycin (BLM)-induced lung fibrosis^11–13^, which, in contrast to the IPF lung, features relatively rapid inflammatory onset, epithelial regeneration, and fibrosis resolution^14,15^.

Although recent single-cell (sc) RNA-seq studies have significantly advanced our understanding of the IPF lung cellular composition^4–7,16–18^, they lack insights into tissue architecture and cellular interplay in a spatial context. Spatially resolved transcriptomics (SRT) enables RNA profiling of intact tissue^19–22^ and can illuminate dynamic cellular interactions in lung tissue^23–25^. However, a transcriptome-wide map of extensive areas of the fibrotic lung is currently missing.

Here, we applied SRT to map the fibrotic lung landscape in human IPF and the BLM mouse model. We integrated SRT with scRNA-seq data to characterize the AbBa cell microenvironment and delineate the dynamic crosstalk between alveolar epithelial cells, myofibroblasts, fibroblasts, and pro-fibrotic macrophages. These first-of-its-kind spatial atlases broaden our understanding of the IPF cellular interplay and unveil key convergent and divergent pathways in human IPF and the BLM mouse model.

## Results

### Spatial transcriptomics of healthy and IPF lungs

We generated transcriptome-wide spatial profiles of freshly frozen human lung biopsies from four IPF patients (IPF 1-4, collected during lung transplantation) and four subjects with no known lung disease (healthy controls; HC 1-4, “B0”, collected post-mortem) using the Visium Spatial Gene Expression platform (**Fig. 1a,b**). For each IPF patient, three biopsies (“B1”, “B2”, “B3”) reflecting increasing extent of fibrotic injury within the same donor were selected (**Fig. 1a)**.

**Figure 1.**
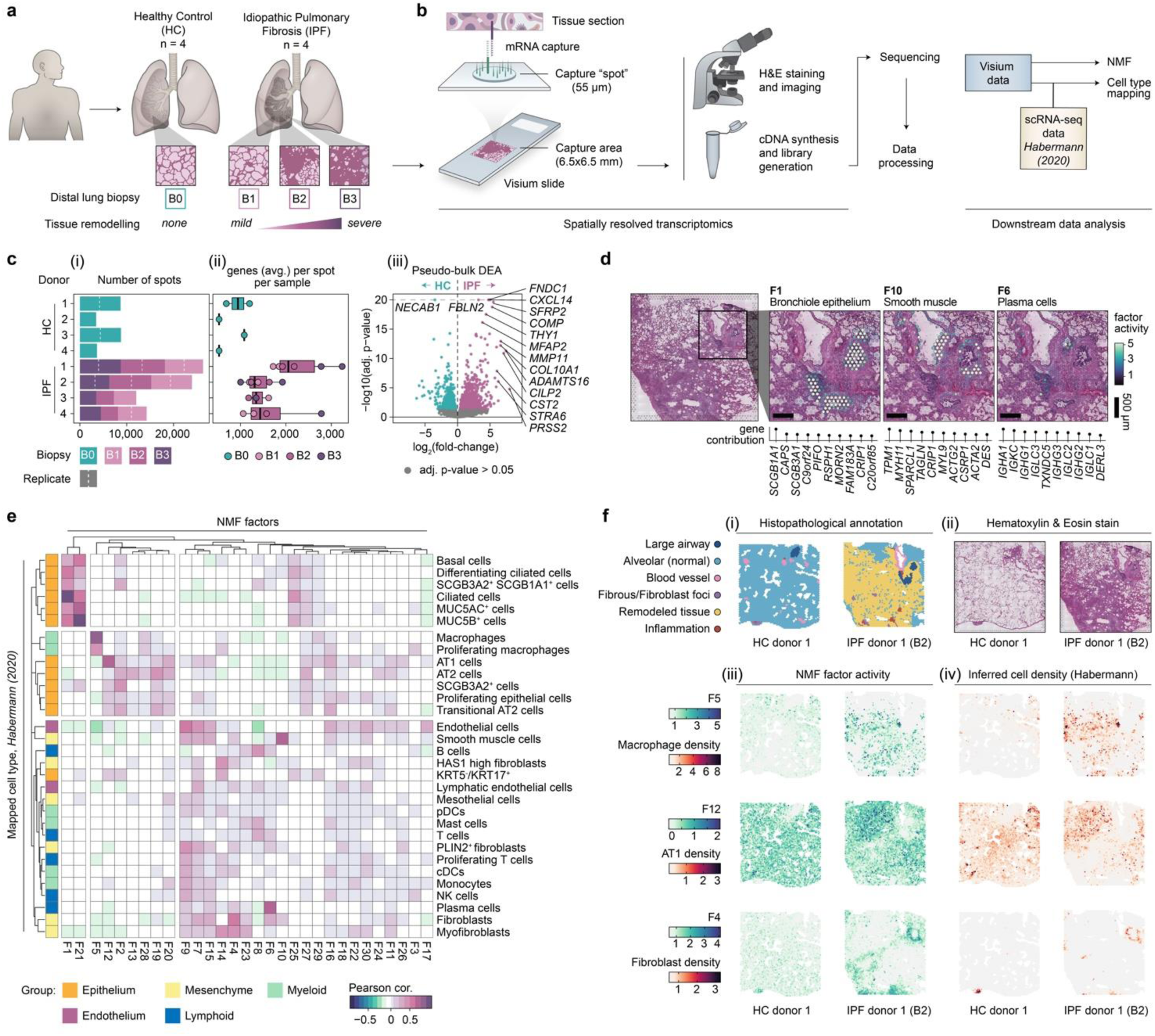
Spatial transcriptomic profiling of human pulmonary fibrosis. **a)** Tissue sections from distal lung biopsies from healthy controls (HC; B0; n=4) and IPF patients (n=4), were sectioned and analyzed using the Visium Spatial Gene Expression technology. Three biopsies exhibiting progressive tissue remodeling (B1-3) were selected from each IPF donor. **b)** Schematic illustration of the Visium workflow and subsequent data processing steps. NMF was used for dimensionality reduction, generating 30 distinct factors. Cell type distributions were inferred through integration with a scRNA-seq dataset published by Habermann et al. (2020; GSE135893). **c)** Summarizing descriptions of the data, including the number of Visium capture spots per sample (i), the average number of unique gene detected per spot (ii), and a pseudo-bulk differential expression analysis (DEA) comparing pooled HC and IPF Visium data per donor to identify significant differentially expressed genes between condition based on data from entire tissue sections (iii). **d)** Spatial distribution maps for selected NMF factors that correspond histological and/or transcriptional profiles of bronchiole epithelium (F1), smooth muscle (F10), and plasma cells (F6). **e)** Correlation (Pearson) heatmap of NMF factor activity and inferred cell type densities, using the Habermann et al. scRNA-seq data set, across spots. **f)** Histopathological annotations performed on sections from each HC and IPF biopsy (i) based on the H&E stained Visium sections (ii). Visualizing spatial NMF activity (iii) and inferred cell type densities (iv), confirms the co-localization of highly correlated factor-cell pairs. H&E, hematoxylin and eosin; NMF, non-negative matrix factorization; DEA, differential expression analysis; AT1, alveolar type 1 cells; AT2, alveolar type 2 cells; pDC/cDCs, plasmacytoid/classical dendritic cells; NK cells, natural killer cells.

We analyzed an average of around 4000 spots (each spot representing a transcriptome of the tissue covering the spot) per tissue section (**Fig. 1c (i)**), capturing an average of >1500 unique genes per spot (**Fig. 1c (ii)**). We observed a higher average number of genes per spot and transcript count levels in IPF samples compared to HC, likely due to disease-associated differences in cellular density between the samples. A pseudo-bulk differential expression analysis (DEA) between IPF and HC samples identified a total of 1469 differentially expressed genes (DEGs) (**Fig. 1c (iii)**), including genes associated with fibroblasts, previously reported to be upregulated in IPF (*FNDC1*, *COL10A1, THY1*)^26^, as well as matrix metalloproteinases (MMPs)^27^ and genes involved in IPF-associated signalling pathways (*SFRP2*, *WNT10A*, *TGFBI*)^28,29^. Many of the upregulated genes in IPF samples mapped to areas of remodelled tissue.

#### Deconvolution of spatial gene expression identifies morphological structures and cell types

The data were deconvoluted into 30 “factors” using non-negative matrix factorization (NMF)^30^ (**Fig. 1d**). These factors revealed gene signatures of distinct cell types and structures including mixed bronchiolar epithelial cell types (Factor 1; F1), smooth muscle cells (F10), and plasma and B cells (F6). The spatial distribution of cell-type densities was further inferred by integration^31^ with an IPF-derived scRNA-seq dataset^5^ (referred to as “Habermann (2020)”). This revealed a distinct group (F1 and F21) that correlated with ciliated airway cell types, including basal cells, club cells, ciliated cells, and MUC5B+ cells (**Fig. 1e**), in line with spatial mapping of F1 activity to bronchial epithelium. Other factors correlated specifically with the alveolar compartment, including alveolar macrophages (spatial overlap with F5; **Fig. 1f**), AT1 cells (spatial overlap with F12 and annotated alveolar tissue; **Fig. 1f**), and AT2 cells. An additional group of factors corresponded to immune cells and stromal components of the lung, including lymphocytes, endothelial cells and fibroblasts (spatial overlap with F4 and areas labelled as fibrous or remodeled tissue; **Fig. 1f**). Several factors could not be clearly attributed to specific cell types/groups, likely representing a more complex mixture of cells, cell types not annotated in the reference dataset, and/or novel cell states. This included F16 in the alveolar compartment of HC and IPF lungs (**Fig. 1e)**, dominated by prostaglandin signaling genes and AT1, AT2, and fibroblast marker genes.

#### Dissecting factor activity reveals pathways and cellular interactions

Further examination of factor distribution across samples revealed 11 factors that were more prevalent in IPF compared to HC (**Fig. 2a**). These factors associated with important IPF cell morphologies/processes (**Fig. 2b**) including ECM-related pathways, and overlapped with regions of tissue fibrosis (F4 and F14) or classic IPF “honeycomb” formations (F5 and F21), whereby F5 displayed markers of dendritic cells and macrophages, whilst F21 presented a *MUC5B*-expressing airway epithelial signature. The F21 profile might reflect a previously identified MUC5b+, BPIFB1+, SCGB3A1+ IPF-associated cell population^32^. In line with our SRT data, MUC5b expression has previously been localized to honeycomb cysts, and *MUC5B* polymorphisms have been linked to IPF risk^33^. F9 appeared to overlap with alveolar regions in IPF tissues and was dominated by genes related to oxidative stress, inflammation, ECM remodeling, and vascular changes, indicating early inflammatory and fibrotic responses, or potential protective mechanisms in the non-remodeled tissue.

**Figure 2.**
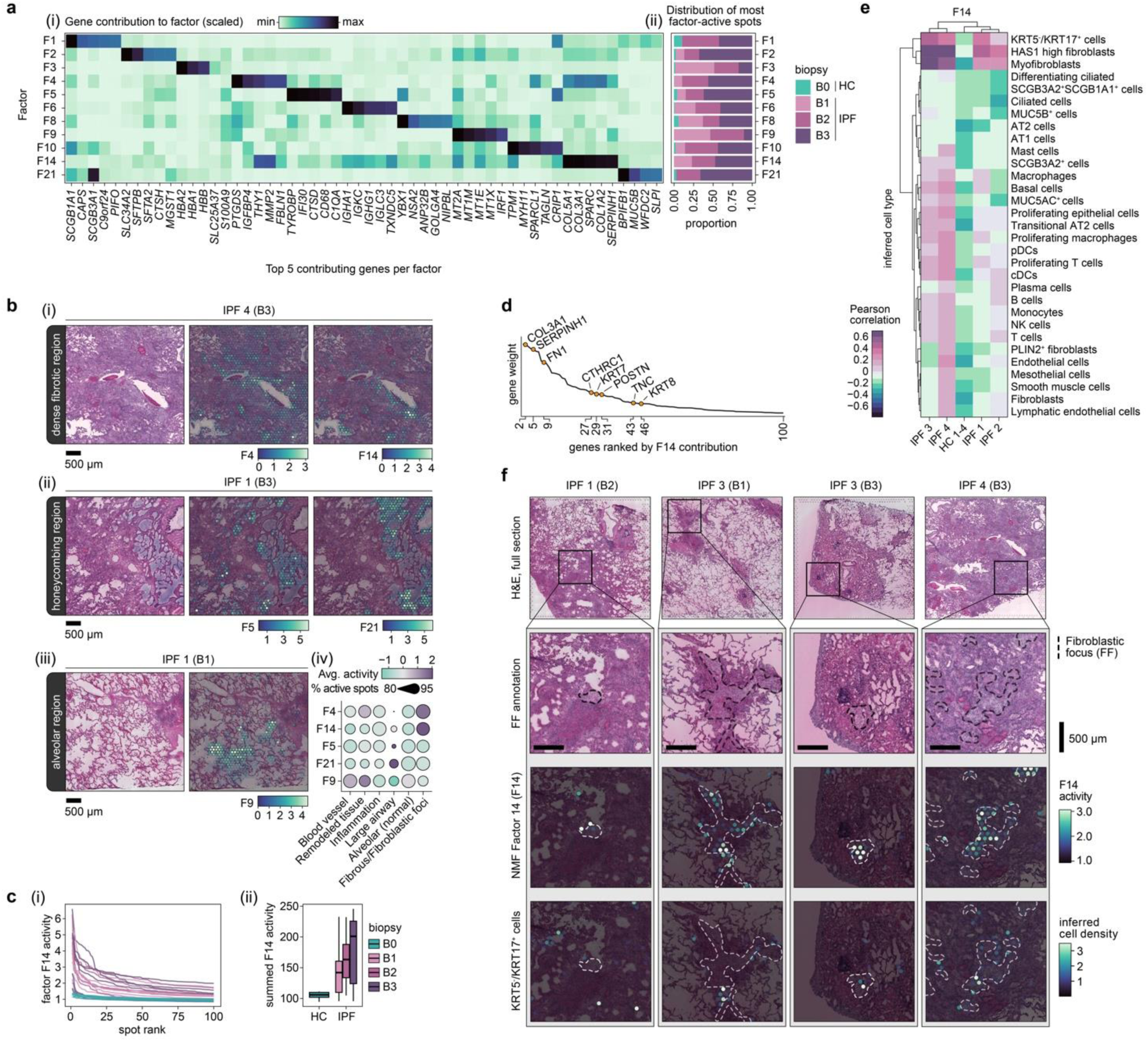
Disease-associated signatures revealed by non-negative matrix factorization. a) NMF identified signatures over-represented in IPF tissue. Their relative contribution to each factor (scaled) is displayed for the top five contributing genes per factor (i), and the proportion of spots with the highest activity (99^th^ percentile) by condition and biopsy severity grade (ii). b) Spatial representation of selected NMF factors across IPF lung sections, demonstrating distinct localization patterns. F4 and F14 marked heavily fibrotic regions (i), F5 and F21 associated with honeycombing structures (ii), and F9 were seen in alveolar regions (iii). Displaying the average activity (scaled and centred) and detection rate (percentage of spots with increased activity) within the annotated histological regions across all biopsies (iv) c) Activity profile of the top 100 ranked spots per sample based on F14 activity, highlighting a consistent distinction between HC and IPF tissues (i), further summarized by summing the F14 activity levels displayed in (i) and grouped based on biopsy remodelling extent (B0-3) (ii). d) The contribution ranking of the top 100 genes for F14 based on gene weight (contribution) to the factor, with keratins, collagens, and other fibrosis-related genes emphasized. e) Correlation heatmap between F14 activity and densities of inferred cell types within spatial spots, capturing potential co-localization of F14 and cell types (strong correlation suggests spatial co-occurrence). f) Visualization of fibroblastic focus (FF) annotations, F14 activity, and the distribution of inferred KRT5-/KRT17+ cells, providing a spatially integrated view of the fibrotic niche. NMF, non-negative matrix factorization.

Among the ECM/fibrosis-related factors, F14 was highly active in IPF, particularly in the more severely remodeled tissue (**Fig. 2c**). In addition to various collagens and fibrosis-related genes, keratins such as *KRT7* and *KRT8* also contributed notably to the factor signature (**Fig. 2d**). F14 activity correlated with inferred cell type densities of KRT5-/KRT17+ AbBa cells, myofibroblasts, and the recently described HAS1-hi fibroblast subtype^5^, specifically in the IPF samples (**Fig. 2e**). One lung (IPF donor 2) demonstrated a weaker correlation between F14 and the KRT5-/KRT17+ AbBa cell type, possibly reflecting interindividual heterogeneity in IPF cellular processes. Visual inspection confirmed that F14 positive spots coincided with the correlated cell types, and revealed that F14 activity spatially aligned with fibroblastic foci (FF) (**Fig. 2f**), a histological feature of active tissue remodeling^24,34,35^. Spots with elevated KRT5-/KRT17+ AbBa cell densities were predominantly situated along the FF borders, confirming the previously proposed positioning of these cells within the fibrotic human lung^4^. Importantly, our NMF approach thus identified a signature encompassing the KRT5-/KRT17+ AbBa cell type independently of scRNA-seq data, placed in its spatial histological context across IPF samples.

### Characterization of the aberrant basaloid epithelial and fibrotic niche in IPF

To better understand the cell type heterogeneity in the FF-specific factor, F14, we isolated its most active spots (denoted F14^hi^) and identified five distinct sub-clusters, denoted F14^hi^ C0-C4 (**Fig. 3a**). Defining genes of C0 corresponded to markers of the KRT5-/KRT17+ AbBa cell type (e.g. *PRSS2, KRT7)* ^4,5^, characteristically devoid of the basal cell marker *KRT5*. The remaining four F14^hi^ clusters expressed genes corresponding to fibroblasts/myofibroblasts (C1 and C2), macrophages (C3), and basal and secretory airway epithelial cells (C4). Based on the marker gene profiles, C1 and C2 appeared to represent fibrotic populations with distinct roles, whereby C1 displayed a matrix deposition and scar formation profile, while C2 had markers indicative of stress responses (metallothioneins), immune modulation (*CCL2*, *FCN3*), and vascular interactions (*ENG*, *THBD*), likely reflecting diverse fibroblast phenotypes within the fibrotic niche of IPF lungs. Spatial inspection revealed alignment of C0-spots with the edges of FF (**Fig. 3b**), mirroring the spatial distribution of inferred KRT5-/KTR17+ AbBa cell densities (**Fig. 2f**) and corroborating C0 as a refined AbBa-dense population within F14. In contrast, the cluster displaying fibroblast markers resided within the FF core.

**Figure 3.**
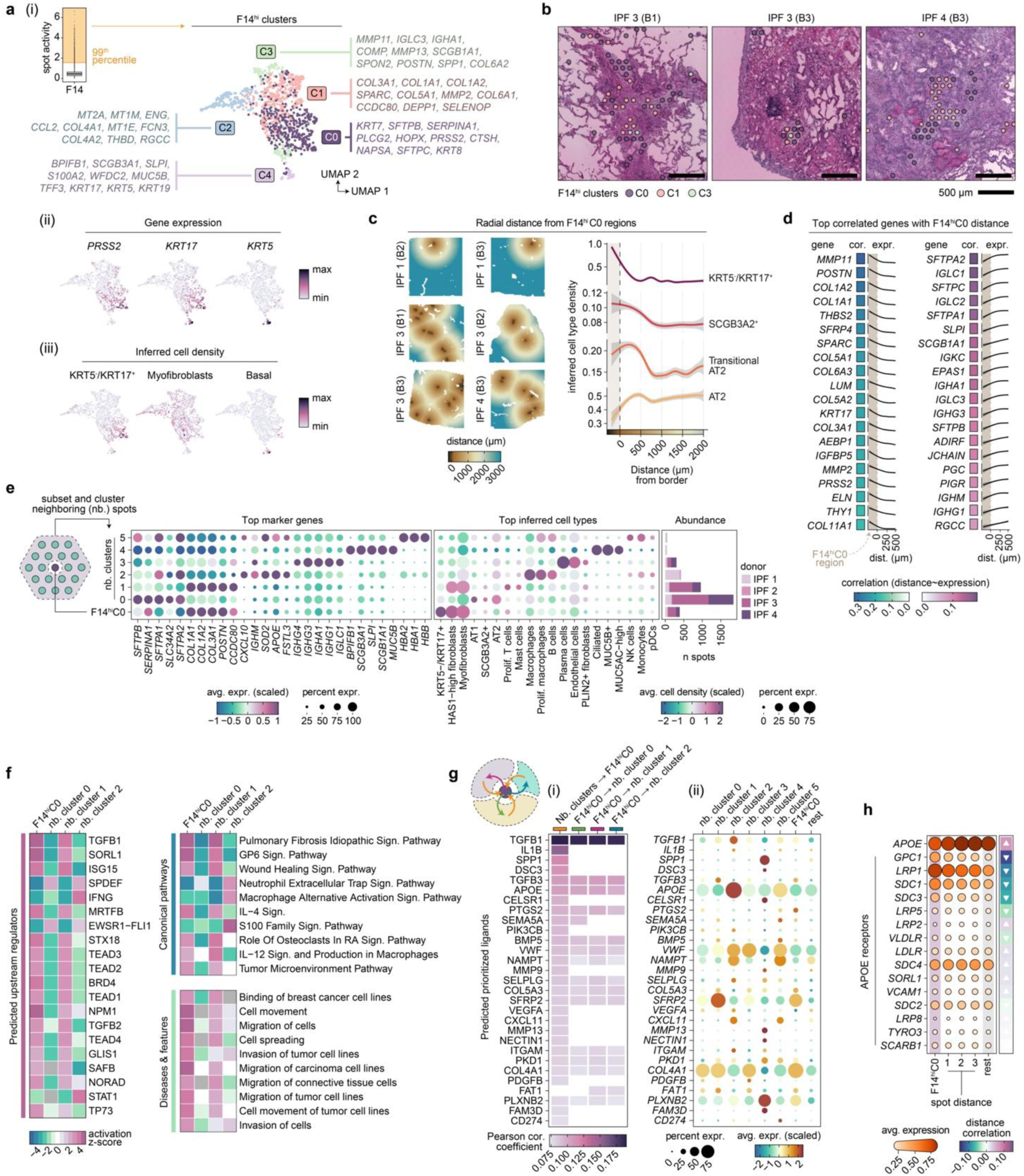
Cellular and molecular deconvolution of the aberrant basaloid niche. **a)** Clustering of the 99^th^ percentile of F14 active (F14^hi^) spots identified five distinct clusters (F14^hi^C0-4), visualized in UMAP space, with the corresponding top ten gene markers listed (i). Expression of the KRT5-/KRT17+ aberrant basaloid cell gene markers PRSS2 and KRT17, along with the absence of KRT5, overlapped with F14^hi^C0 (ii). Inferred cell-type densities highlighted the prominence of KRT5-/KRT17+ cells in F14^hi^C0, while lacking inferred basal cells (iii). **b)** The spatial location of F14^hi^ clusters within FF in representative IPF biopsies. F14^hi^C0 was predominantly observed at the periphery of FF, while F14^hi^C1 (fibroblasts) localized to the FF core. **c)** The spatial tissue location of cluster F14^hi^C0 across IPF lung sections with an analysis of inferred cell type densities relative to the radial distance from the F14^hi^C0 boundary (distance = 0). Distances below zero corresponds to spots within F14^hi^C0. Smoothed cell type densities produced by fitting a generalized additive model (GAM) to the data, where gray shadings indicate 95% confidence interval **d)** Gene expression correlation analysis within a 500 µm radius from the F14^hi^C0 border identified the top 20 positively and negatively associated genes based on correlation (Pearson) values. **e)** Neighboring (nb.) clusters of the F14^hi^C0 regions were generated by further clustering of spots within a 2-spot distance (∼ 200 µm) of the F14^hi^C0 borders. Dot plots displays the top marker genes of each F14^hi^C0 nb. Cluster and the inferred cell type densities of selected cell types to highlight their abundance within each cluster. Bar chart shows the number of spots per donor labeled with each F14^hi^C0 nb. cluster. **f)** Enrichment analysis in Ingenuity Pathway Analysis (IPA) based on marker genes (adj. p < 0.05) for nb. clusters 0-2 when compared against each other, and F14^hi^C0 markers when compared against all other spots in the IPF samples. Heatmaps of activation z-scores of top 20 significant predicted upstream regulators, and top 10 enriched canonical pathways and diseases and functions. **g)** Cell-cell communication analysis using NicheNet within the F14^hi^C0 niche. Prediction of prioritized ligands acting upon the F14^hi^C0 and nb. clusters 0-2 regions (i), with mean expression levels of the ligands in each cluster to deduce the potential origin of the ligand (ii). **h)** Average expression levels of *APOE* and its canonical receptors within F14^hi^C0 and their change over radial spot distance (3 spots; ∼300 µm), where “rest” corresponds to the background expression observed across all remaining spots across samples. Directional arrows indicating correlation (Pearson) trends based on expression over spot distance. UMAP, uniform manifold approximation and projection; FF, fibroblastic foci.

#### Cellular crosstalk and molecular signaling in the IPF AbBa microenvironment

We found a higher abundance of AT2 cells and transitional AT2 cells around the F14^hi^C0 AbBa niche, compared to more distant regions (**Fig. 3c)**, whereby the peak of transitional AT2 cell density was observed at a shorter distance compared to the peak of AT2 cells. This suggests a possible differentiation lineage from AT2 to transitional AT2 cells and towards AbBa cells, consistent with a previously proposed cell trajectory^5^, captured in space. Additionally, the proximity of SCGB3A2+ secretory cells to F14^hi^C0 spots aligns with previous findings suggesting them as another potential source for AbBa cells ^5,32^.

We observed a decline in matrix remodeling and fibrosis-associated genes (e.g., *MMP11, POSTN, COL1A2)* with increasing distance from F14^hi^C0 (**Fig. 3d**), indicating elevated fibrotic activity around AbBa cells. Conversely, genes linked to alveolar function and immune response (e.g., *SFTPA2, SFTPC*, *SLPI)* showed lower expression within C0 compared to its immediate surroundings. A group of immunoglobulin-related genes (e.g., *IGLC1, IGKC, PIGR*) resided near AbBa cell dense areas, but not within, implying a differential immune response or possible exclusion of certain immune elements from the AbBa microenvironment.

Further analysis of areas neighboring (nb) the F14^hi^C0 spots identified clusters containing alveolar epithelial cells (nb. cluster 0), fibroblasts/myofibroblasts (nb. cluster 1), alveolar macrophages (nb. cluster 2), and plasma cells (nb. cluster 3) (**Fig. 3e**), allowing us to study regulatory molecules and signaling within and between clusters. Upstream regulator and pathway enrichment analyses performed in Ingenuity Pathway Analysis (IPA) predicted upstream activation in F14^hi^C0 and nb. cluster 1 of molecules (including TGFB1, TGFB2, MRTFB, TEAD1-4, ISG15) known to be involved in fibrosis (**Fig. 3f)**. The canonical pro-fibrotic cytokine TGF-β (encoded by *TGFB1* and *TGFB2*) plays a significant role in IPF^28,36^ and has been implicated in ADI cell formation and inhibition of differentiation towards AT1 cells^37^. MRTFB regulates myofibroblast differentiation^38^, whilst TEAD family members (part of YAP/TAZ co-activator complex) are key effectors of profibrotic pathways including Hippo-, TGF-β, and Wnt signaling^39–41^, implicated in tissue regeneration and in fibrosis^29,42^. The p53 modulator, ISG15, implicated in age-related signaling^43^, was a predicted activated upstream regulator of F14^hi^C0. Enrichment of IPF-, glycoprotein VI (GP6)-, and wound healing signaling pathways, along with pathways associated with cell movement and migration, further supports an active fibrogenic node.

Prediction analysis of ligand-target interaction^44^, with directional information preserved (Methods), identified further cell-cell communications within the F14^hi^C0 microenvironment, including TGFB1, IL1B, and SFRP2 (**Fig. 3g**). SFRP2 (a Wnt signaling modulator) expression distinctly originated from the neighboring fibroblast cluster, implicating potential autocrine/paracrine Wnt signaling between (myo)fibroblasts, alveolar epithelial, and AbBa cells. Furthermore, the predicted ligand apolipoprotein E (*APOE*), with its receptor SORL1 being an upstream regulator of the AbBa-dense cluster, was highly expressed in the macrophage cluster, alluding to a monocyte-derived and M2-like profile of the neighboring macrophage population^45,46^. By analyzing the gene expression data of all annotated APOE receptors across the F14^hi^C0 region distance, we identified an inverse expression pattern between *APOE* and several of its receptors (**Fig 3h**). Glypican 1 (*GPC1*), LDL receptor-related protein 1 (*LRP1*), and syndecan 1 (*SDC1*) were more highly expressed within, and in close proximity to, the F14^hi^C0 region. These observations suggest a potentially under-recognized role for apolipoprotein signaling within the AbBa cell fibrotic niche in IPF.

### Spatially resolved transcriptomics in a mouse model of pulmonary fibrosis

To increase understanding of the translational predictivity of the BLM mouse model for human IPF, we generated SRT data from mouse lung samples collected at day 7 (d7) and day 21 (d21) following BLM or saline (vehicle) administration (**Fig. 4a**).

**Figure 4.**
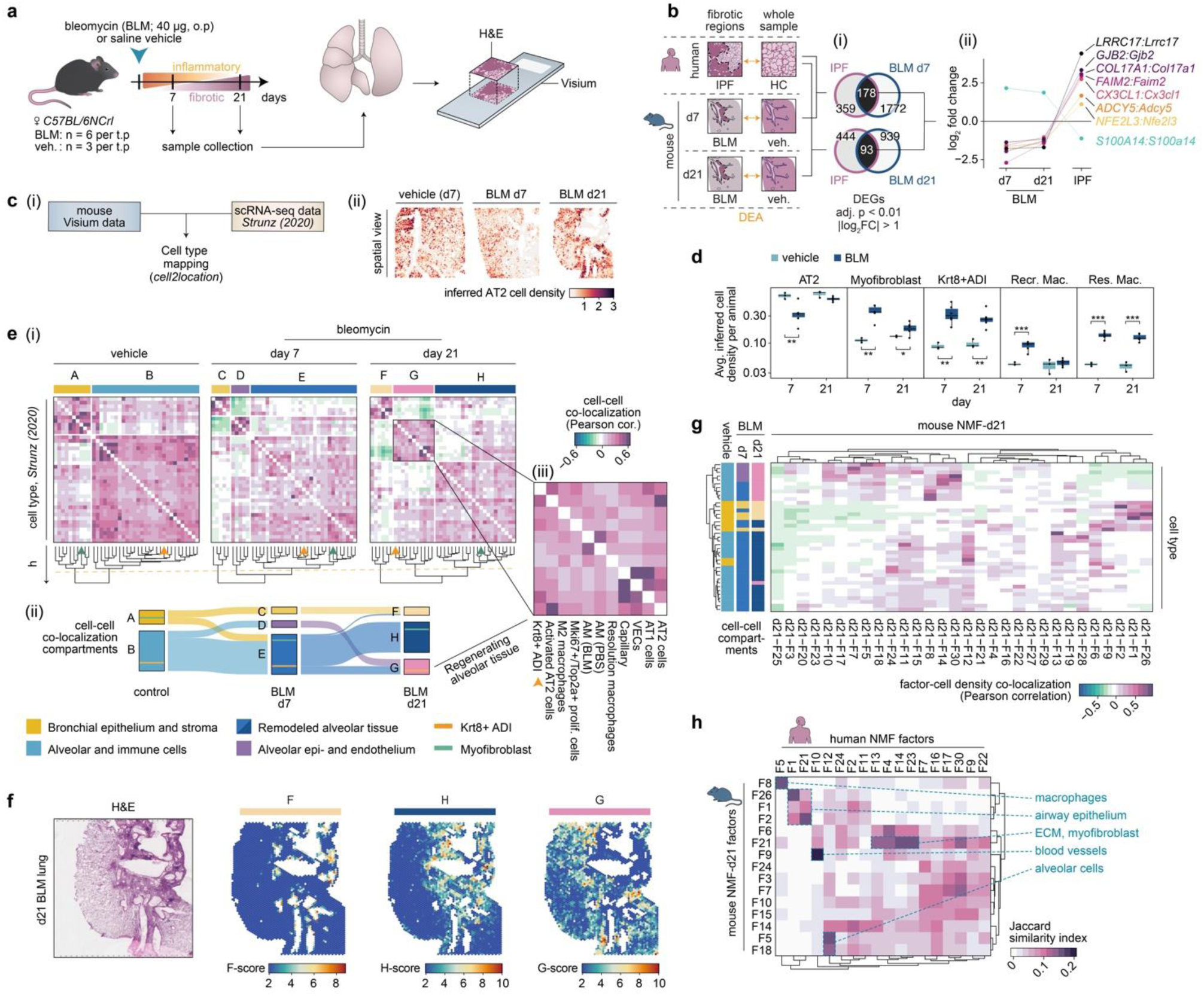
Comparative spatial analysis of pulmonary fibrosis in mouse and human. **a)** Experimental design for the mouse bleomycin (BLM) lung injury model. Mouse lungs were collected at days 7 (d7) and 21 (d21) post BLM or vehicle administration for spatial transcriptomic analysis using the Visium platform (n=6 BLM, n=3 vehicle per time point). **b)** Differential expression analysis (DEA) compared annotated fibrotic regions in human IPF and BLM-treated mouse tissue against respective controls. Venn diagrams of differentially expressed genes (DEGs) specific and shared between human IPF and mouse d7 or d21 post-BLM treatment (i), where the shared genes exhibiting inverse expression patterns between human IPF and the BLM samples are further explored (ii). **c)** Schematic overview of the scRNA-seq data integration, using Visium data and the annotated scRNA-seq data set published by Strunz et al. (2020; GSE141259) as input for cell2location to infer spot cell type densities (i). Exemplified by the inferred AT2 cell density in selected Visium samples across time points (ii). **d)** Averaged cell type abundance per animal, comparing densities between timepoints and treatments for selected cell types. Welch Two Sample t-test (two-sided; nVeh. = 3, nBLM = 6, per time point) was used to test for significance between groups, * p < 0.05, ** p < 0.01, *** p < 0.001. Center line, median; box limits, upper and lower quartiles; whiskers, 1.5x interquartile range; points, value per animal. **e)** Inferred cell-cell correlation heatmaps display distinct cellular co-localization compartments that changes across condition and time. Compartments (A-H) were identified based on the same height, *h*, cutoff (orange dashed line) (*h* = 1.5) of the hierarchical clustering for the selected data subsets (i). Sankey diagram depicting the shift in cell types within each compartment from vehicle controls to BLM d7 and then BLM d21, illustrating the cellular spatiotemporal dynamics within fibrotic mouse lungs (ii), with Krt8+ADI (orange line) and myofibroblast (green line) populations highlighted how they move across compartments. Zooming into compartment G (iii), Krt8+ ADI cells (orange arrow) are found co-localizing with alveolar epithelial cells and macrophages. **f)** Computed scores for each compartment (F-H scores), calculated by summing the cell type densities of the compartment-associated cell types, displayed in a BLM d21 lung section alongside the H&E staining of the same section. **g)** NMF was performed on the d21 subset and the factor activities in each spot were compared with inferred cell-type densities using Pearson correlation and hierarchical ordering. The cell type group colors correspond to their respective compartments (A-H) based on the prior analysis and highlight sets of factors that strongly matches distinct or groups of cell types, largely capturing a similar BLM d21 compartmentalization. **h)** Comparison of gene contributors to the human derived NMF analysis with the mouse d21 NMF analysis, by computing and visualizing the Jaccard similarity coefficient based on the top 100 genes contributing to each factor. Heatmap displays filtered results based on factors having a Jaccard index of > 0.1 with at least one other factor, to exclude factors with no apparent overlap between species. NMF, non-negative matrix factorization; DEA, differential expression analysis; DEG, differentially expressed genes; ADI, alveolar differentiation intermediate; AM, alveolar macrophages; Recr. Mac., recruited macrophages; Res. Mac., resolution macrophages; VECs, vascular endothelial cells; ECM, extracellular matrix.

Healthy alveolar regions accounted for 80-90% and 30-50% of the total number of spots in saline and BLM challenged lungs, respectively. Remaining spots in the BLM challenged samples were labelled as areas of tissue damage or remodeling. A pseudo-bulk DEA between BLM and vehicle controls identified a total of 3214 and 3787 DEGs at d7 and d21, respectively.

#### Comparative analysis of gene expression and cellular composition

We identified differentially expressed genes (DEGs) in annotated fibrotic areas compared to control samples in the mouse model and analyzed their overlap with DEGs in IPF (**Fig. 4b)**. Numerous DEGs overlapped between mouse and human (178 between IPF and d7 BLM fibrotic regions, and 93 between IPF and d21 BLM), with eight DEGs displaying contrasting fold-change directionality. Among the latter, most are involved in ECM organization (*COL17A1*^47^), inflammatory signaling (*CX3CL1*^48^), and apoptosis regulation and cellular adhesion (*S100A14*^49^, *FAIM2*^50^).

While these genes may play a role in fibrosis in both conditions, Their inverse expression patterns of these DEGs suggest divergent roles in human IPF compared to the mouse BLM model.

Cell type deconvolution was performed using a lung scRNA-seq dataset generated in the BLM mouse model^11^ (referred to as “Strunz (2020)”). Spatial visualization of cell type densities demonstrated accurate mapping to relevant tissue regions, where alveolar epithelial cells were inferred in healthy alveolar tissue (**Fig. 4c**).

Pronounced differences in cell type densities were observed between BLM and vehicle groups, including resolution (M2 polarized) macrophages and Krt8+ADI cells (**Fig. 4d**). AT2 cell abundance decreased at d7 but showed recovery by d21. The apparent influx of recruited (pro-inflammatory) macrophages at d7 normalized by d21, confirming resolution of acute inflammation.

#### Spatial compartmentalization reveals dynamic lung tissue remodeling in response to BLM

Co-localization analysis revealed dynamic spatial compartmentalization of cell types within spots, capturing the spatiotemporal dynamics of fibrogenesis and indicating lung tissue remodeling in response to BLM injury (**Fig. 4e**). In vehicle control lungs, we identified two compartments consisting of bronchial epithelial (A) and alveolar (B) tissue, outlining the uninjured lung architecture. In the d7 BLM-challenged lungs, prominent cell densities consisted of bronchial epithelium (C), alveolar epithelium and alveolar capillary endothelium (D), and remodeled alveolar tissue marked by fibroblasts and myofibroblasts (E). At d21 the cellular composition of the compartments was altered, so that in addition to bronchial epithelium (F) and fibrotic, remodeled alveolar tissue (H), we observed a compartment (G) characterized by alveolar epithelium macrophages and Krt8+ADI cells, exhibiting a profile of regenerating alveolar tissue (**Fig. 4e**). Spatial mapping confirmed that F aligned with bronchial structures, H coincided with fibrotic/remodeled tissue, while G was present along the borders of fibrotic areas and extending into intact tissue (**Fig. 4f**).

#### Comparative analysis of regenerative signatures reveals divergent epithelial responses

Given that day 21 in the mouse model reflected an established stage of fibrosis with minimal acute inflammation, we focused on this time point for comparison with IPF. NMF application to the mouse d21 data (mmNMF_d21_) generated 30 factors. Factor activity and cell type abundance co-localization analysis largely reflected the d21 BLM compartmentalization, affirming that NMF effectively captures patterns comparable to the cell type deconvolution approach (**Fig. 4g**). The regenerating alveolar epithelial compartment (G) was represented by a set of factors primarily reflecting AT2 cells (F30), alveolar and resolution macrophages (F8), or activated AT2 and Krt8+ADI cells (F14). Factors F18, F5, and F7 predominantly represented AT1 and endothelial cells.

We further compared mmNMF_d21_ factors with factors identified by IPF NMF analysis (hsNMF) **(Fig. 4h)**. The top contributing genes showed an overall weak overlap between human and mouse factors. However, factors associated with distinct morphological features, such as smooth muscle cells (SMC), blood vessels, and ciliated airway epithelium, demonstrated more pronounced overlap, highlighting conserved signatures normal lung structures, compared to disease or injury responses. Notably, factors containing transcriptional signatures for the human KRT5-/KRT17+ AbBa cells (hsNMF-F14) and mouse Krt8+ADI cells (mmNMF_d21_-F14) had a limited overlap.

### Translation of the fibrotic microenvironment

#### Contrasting fibrogenic and regenerative responses in human IPF and BLM-induced lung fibrosis in mouse

We analyzed the spatial correlation of the factors containing KRT5-/KRT17+ AbBa (hsNMF-F14) in human IPF samples and the factors containing Krt8+ADI cells (mmNMF_d21_-F14) in the mouse BLM samples. HsNMF-F14 activity predominantly correlated with fibroblasts (HAS1-hi), myofibroblasts, and KRT5-/KRT17+ AbBa cells. Conversely, mmNMF_d21_-F14 activity primarily correlated with Krt8+ADI cells and AT2 cells, while showing a weaker correlation with myofibroblasts. Additionally, mmNMF_d21_-F14 showed correlation (albeit weaker) with AT1 cells, unlike hsNMF-F14 **(Fig. 5a)**, in line with the distinct fibrogenic environment in the human aberrant basaloid niche.

**Figure 5.**
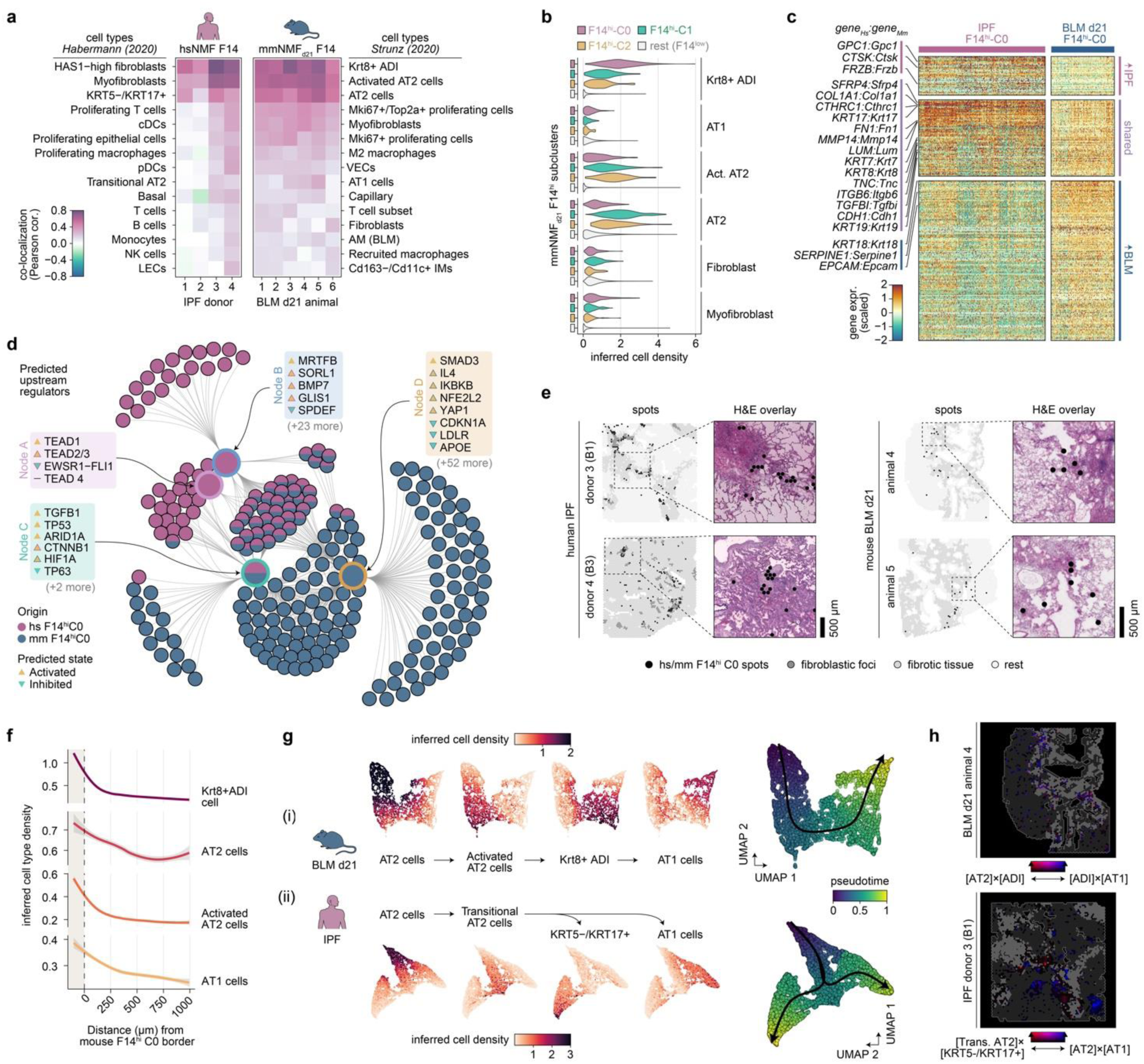
Translational dissection of the fibrotic niche and cellular dynamics. **a)** Correlated activity, per individual, of the human (hs) NMF-F14^hi^ and mouse (mm) NMF-F14^hi^ factors with the 15 highest correlated inferred cell types using the Habermann (human) and Strunz (mouse) scRNA-seq data sets. **b)** Distribution of selected cell densities within subclustered mmNMFd21-F14^hi^ spots, demonstrating high abundance of Krt8+ ADI cells within mmNMFd21-F14^hi^C0 while high AT2 cell abundance in mmNMFd21-F14^hi^ C1 and C2. **c)** Integrated IPF and BLM-d21 data sets by converting orthologous gene names, allowed identification of marker genes for hsNMF-F14^hi^C0 and mmNMFd21-F14^hi^C0. Heatmap with scaled and centered marker gene expression, grouped based on whether significant marker genes were elevated in IPF samples, shared, or higher in mouse BLM tissues. A total of 74 genes were found to be shared, while 39 were seen significant in the IPF and 157 in the d21 BLM cluster (adj. p < 0.01, avg. log2FC > 0). **d)** Comparative network plot showing the most significant regulators (p value < 10^−7^, right-tailed Fisher’s exact test) based on IPA upstream analyses of marker genes (adj. p < 0.05) from hsNMF-F14^hi^C0 and mmNMFd21-F14^hi^C0. Inner nodes illustrate groups of regulators sharing genetic influences, and outer nodes represent contributing marker genes. **e)** Spatial mapping of the hsNMF-F14^hi^C0 and mmNMFd21-F14^hi^C0 spots within the tissue sections illustrating the relationship with fibrotic regions, providing a visual correlation with areas of disease pathology. **f)** Radial distribution line graphs for inferred cell densities around the mmNMFd21-F14^hi^C0 niche mapped out a gradient of alveolar cell composition. Smoothed lines produced using local polynomial regression fitting (“loess”), where gray shading corresponds to 95% confidence interval. **g)** Spatial trajectory analysis was carried out by selecting spots containing high inferred densities of the selected cell types AT2, activated or transitional AT2, Krt8+ADI or KT5-/KRT17+, and AT1 cells. Trajectories and pseudotime were thereafter inferred using the Slingshot methodology based on UMAP embeddings of the cell type densities for the selected spots, where AT2 cells were defined as the starting cluster. Deviating trajectories were seen between IPF and BLM-induced lung fibrosis, where in the BLM mouse (i) a single trajectory was observed along the proposed AT2-Krt8+ADI-AT1 lineage, while two divergent trajectories were seen in human IPF (ii) in which aberrant basaloid KRT5-/KRT17+ cells were spatially disconnected from AT1 cells. **h)** Spatial co-localization visualization of the AT2-to-Krt8+ADI (red) and ADI-to-AT1 (blue) inferred cell densities in mouse and the transitional AT2-to-KRT5-/KRT17+ (red) and AT2-to-AT1 (blue) densities in human, by computing cell density products and visualizing their intensities along a red-blue axis. A mixture of the density products will appear purple, and the brightness corresponds to value intensity (black = 0). Spots with values of zero are excluded. Tissue outlines and areas of fibrosis (gray) illustrated for guidance. In mouse, signals along the entire AT2–AT1 trajectory is found mixed near borders of fibrosis, while in human, spatially isolated regions display high co-localization intensities of cells from the two inferred trajectories. ADI, alveolar differentiation intermediate; IPA, Ingenuity Pathway Analysis.

To further compare the gene signatures of the AbBa (IPF) or ADI (BLM) niches, we refined mmNMF_d21_-F14 and clustered the spots based on gene expression (Methods), and identified four clusters (mmNMF_d21_-F14^hi^ C0-3), where cluster 0 (mmNMF_d21_-F14^hi^ C0) exhibited the strongest association with Krt8+ADI cells **(Fig. 5b)**. We detected shared marker genes between hsNMF-F14^hi^ C0 and mmNMF_d21_-F14^hi^ C0, including several collagens and ECM-related genes (e.g., *COL1A1, FN1*, *TNC, CTHRC1*), epithelial cell markers (*CDH1*), and markers for human AbBa cells (*KRT17*) and mouse ADI cells (*KRT8*) (**Fig. 5c)**. This suggests shared traits between AbBa and ADI regions involving ECM remodeling and a basaloid phenotype, further supported by pathway analysis. However, the ADI-related gene signature observed in mouse predominantly engaged pathways related to inflammation and repair, whereas the AbBA-related gene signature observed in human IPF reflected the chronic and progressive nature of IPF, dominated by immune responses and pathways governing long-term tissue remodeling.

For better understanding of the aberrant fibrotic niche drivers, we performed an upstream regulator analysis for hsNMF-F14^hi^ C0 and mmNMF_d21_-F14^hi^ C0 (**Fig. 5d**). Both groups had predicted activation of TGFB1, TP53, and SMAD3, suggesting a conserved TGF-β-related mechanism^28,51,52^ and cellular senescence^53^. hsNMF-F14^hi^ C0-specific regulators included the anti-fibrotic growth factor BMP7, the ApoE receptor SORL1, and GLIS1, a component of the Notch signaling pathway. MmNMF_d21_-F14^hi^ C0 showed activation of oxidative stress and inflammation regulators including as HIF1A, IL4, YAP1, and NFE2L2 (NRF2). Contrary to our previous findings in the human samples suggesting a role for apolipoprotein signaling acting upon hsNMF-F14^hi^ C0 (**Fig. 3f-h**), APOE and its receptor LDLR were predicted as inhibited regulators for the mouse mmNMF_d21_-F14^hi^ C0 spots in this analysis.

Next, we examined the histological context of the mmNMF_d21_-F14^hi^ C0 cluster and found it primarily situated at the junction between healthy and fibrotic tissue, comparable to the localization along the FF border seen for hsNMF-F14^hi^ C0 in the IPF samples. Placement at the remodeling tissue interface supports a transitional niche role for these clusters **(Fig. 5e)**.

The BLM Krt8+ADI transitional cell population is predicted to originate from either AT2 cells or club cells and differentiate into AT1 cells^11^. By assessing the cell type densities in relation to their radial distance from the borders of the mmNMF_d21_-F14^hi^ C0 niche, we identified high densities of AT2 cells, activated AT2 cells, and AT1 cells close to the mmNMF_d21_-F14^hi^ C0 niche (**Fig. 5f**). These observations shared similarities with the corresponding IPF hsNMF-F14^hi^ C0 analysis (**Fig. 3c**), highlighting the absence of AT1 cells around the human AbBa niche.

A spatial trajectory analysis of the alveolar epithelial cell types/states (**Fig. 5g**; ‘Methods’) identified a single spatial trajectory from AT2 cells to activated AT2 cells, ADI cells, and culminating in AT1 cells, in the BLM mouse data. In contrast, the cell composition in the IPF lungs displayed a branching trajectory from AT2 cells through transitional AT2 cells, subsequently diverging into either KRT5-/KRT17+ AbBa or AT1 cells.

Visualizing these trajectories spatially, we observed a separation between transitional AT2–AbBa (fibrosis-associated) and AT2–AT1 (alveoli-associated) niches (**Fig. 5h**).

#### Uncovering immune cell dynamics in pulmonary fibrosis

Our NMF analysis revealed factors in the IPF and the d7 and d21 BLM datasets that shared key marker genes indicative of macrophages (*SPP1*, *CD68*, *APOE*). SPP1+ profibrotic macrophages, displaying an M2 polarization phenotype, have previously been implicated in ECM remodeling and fibrosis development^6,54^. The activity of the selected macrophage factors was largely localized near fibrosis-associated bronchial regions.

A shared histological feature between the IPF lungs and the mouse BLM-injured lungs was the presence of dense immune cell infiltrates embedded within the fibrotic tissue (**Fig. 6a**). In the timepoint-separated mouse NMF analyses, mmNMF_d7_ and mmNMF_d21_, we identified factors prevalent in regions of immune infiltrates. In the human NMF, a similar histological feature was not consistently detected, therefore more targeted donor-specific NMF analyses were performed. Factors in three of the donors (IPF 1-3) were seen to overlap spatially with the observed immune infiltrates. A closer examination of the gene contributions (**Fig. 6b**) and inferred cell type composition (**Fig. 6c**) within these immune-dense regions revealed notable differences. In the BLM mouse model, enrichment of genes such as Cd74 and Coro1a indicated presence of antigen-presenting cells and lymphocytes^55^. Additionally, a distinct factor was identified in proximity to the d21 BLM immune-dense structures (**Fig. 6a**), characterized by a plasma cell signature strongly driven by expression of the IgA heavy chain (*Igha*). Overall, the BLM regions demonstrated a relatively balanced mixture of B, T, and dendritic cells, in contrast to the human IPF samples which showed a pronounced expression of chemokine CXCL13, suggesting a B cell-driven immune mechanism^56^. Indeed, B cells were found to dominate, with T cells and plasma cells playing a lesser role, within the lymphocyte-dense regions in IPF lungs. In accordance with recent descriptions of their presence in both healthy and diseased lungs^57^, lymphocytes and plasma cells may have a modulatory role in the progression of fibrosis.

**Figure 6.**
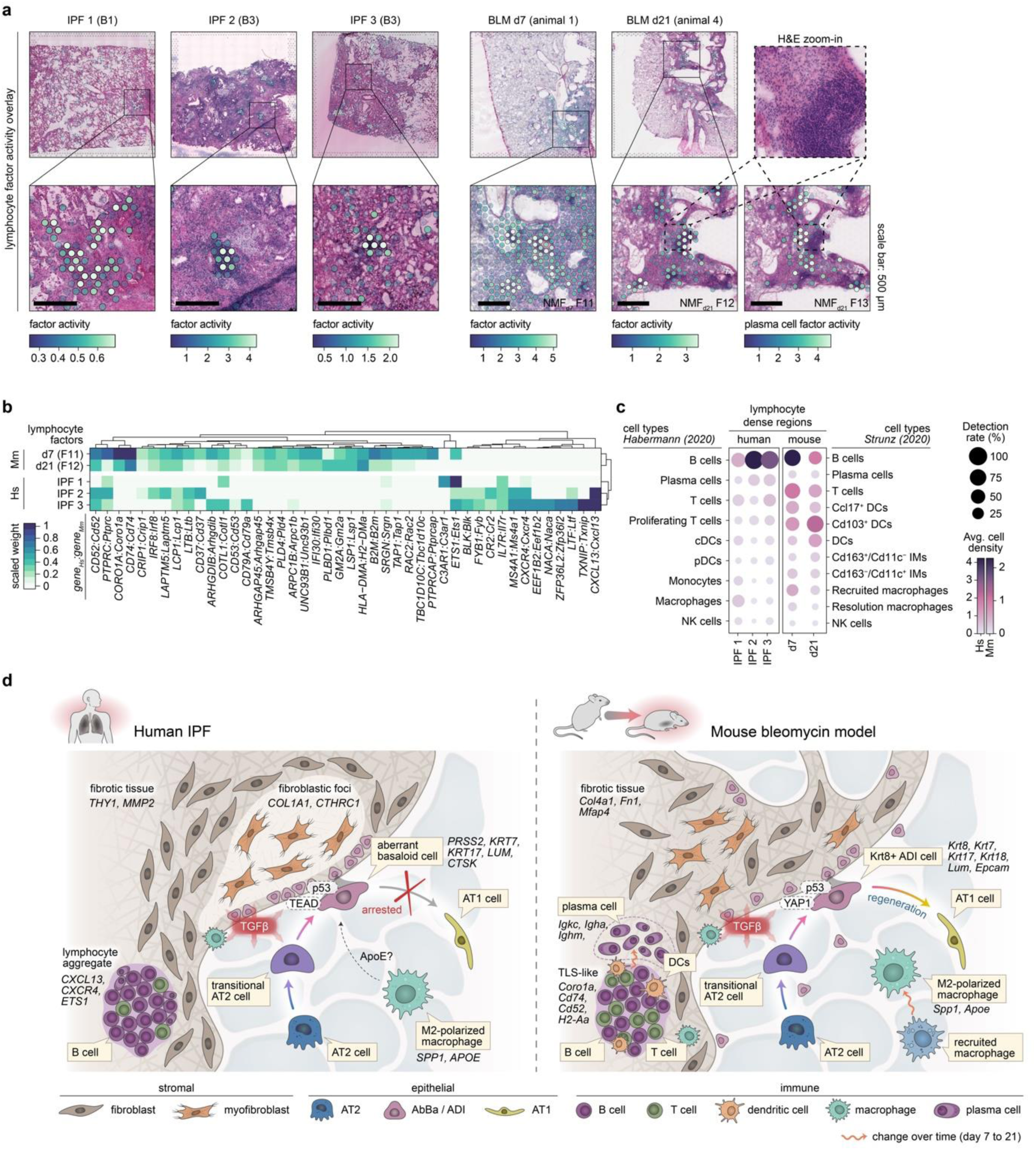
Immune cell signatures and comparative overview of fibrotic mechanisms in human IPF and the bleomycin mouse model. **a)** Spatial visualization of NMF factors overlapping dense lymphocyte / immune cell aggregates in selected human and mouse samples. Scale bars: 500 µm. Imaged at 20X magnification. **b)** Heatmap displays the top contributing factor genes across condition, filtered to show genes with a summed scaled weight above 0.5 across the groups. **c)** Dot plot with inferred cell type densities, for selected immune cell types from the Habermann (2020) and Strunz (2020) data sets, in the most active spots of the selected human and mouse factors. **d)** Schematic summary of the fibrotic niche in human IPF lungs and in mouse BLM-injured lungs, illustrating the proposed cellular interplay within the fibrotic lungs. A key distinction between IPF and the BLM mouse model was centered around the diverging regenerative properties of the IPF-associated AbBa cells versus the mouse Krt8+ ADI cells. While both populations exhibit signs of senescence (p53), the mouse ADI state appears to maintain a functional balance that still prompts it to differentiate into AT1 cells. TGF-beta and Wnt-related (TEAD, YAP1) signaling pathways were central within the fibrotic niche, and the presence of immune cells in proximity to, or within, the severely remodeled tissue implies active fibrogenic modulatory roles. Pro-fibrotic M2-polarized (“resolution”) macrophages with similar gene signatures, expressing *SPP1* (*Spp1*) and *APOE* (*Apoe*), were detected in both human IPF and mouse BLM-injured lungs. In contrast to human IPF AbBa regions, a predicted negative APOE upstream signaling was identified in mouse ADI regions. In mouse, the recruited pro-inflammatory macrophages seen at the early timepoint post BLM-installation were absent by day 21. Establishment of plasma cells adjacent to TLS-like areas in the BLM-injured mice occurred at the later timepoint. AbBa, aberrant basaloid; ADI, alveolar differentiation intermediate; DCs, dendritic cells; IMs, interstitial macrophages; NK cells, natural killer cells; TLS, tertiary lymphoid structure.

Taken together, our analyses delineate distinct cellular trajectories and molecular mechanisms in the fibrotic niche of human IPF and the BLM mouse model (**Fig. 6d**). We highlight the arrested alveolar cell regeneration in IPF versus the active repair in the BLM model, alongside distinct signaling molecules such as TGF-β, ApoE, YAP1, and TEAD, and differences in immune cell presence. These comparative insights underscore the unique aspects of fibrosis in human IPF.

## Discussion

Our study presents a comprehensive comparative genome-wide spatial transcriptome map of the diverse cellular ecosystems and distinct molecular signatures in the human IPF lung and the BLM mouse model.

We propose a central involvement of TGF-β signaling in IPF, alongside other mediators such as TP53, SMAD3, BMP7, MRTFB, TEAD, GLIS1, and APOE, which are linked to senescence, myofibroblast activation and differentiation, Notch and Wnt signaling, apoptosis, and cell migration.

Using data factorization, we identified KRT5-/KRT17+ AbBa and Krt8+ ADI cell populations and their proximate neighborhoods, delineating a critical region within the fibrotic landscape. The complex cross-directional signaling network illustrated within the AbBa niche suggests these cells serve as a transitional core in the IPF lung, whereby AbBa cells orchestrate the fibrotic response, signaling to neighboring cells and modulating the local microenvironment.

TGF-β, a pro-fibrotic cytokine with a significant role in IPF pathogenesis^28,36^, and SMAD3, integral to the TGF-β signaling pathway^58^, were predicted as upstream regulators in human AbBa and mouse ADI fibrotic niches, pointing to a shared TGF-β-driven fibrotic signaling pathway. Furthermore, our data suggest that the role of APOE signaling within the IPF fibrotic niche is more substantial than previously appreciated. This warrants further exploration into the potential regulatory function of APOE in IPF, given its well-documented function in lipid metabolism and its emerging role in immunomodulation and fibrosis^59,60^.

Through tracing the alveolar epithelial cell spatial trajectory, we observe a AT2-ADI-AT1 lineage in the mouse model that is preserved in situ, supporting previous single cell and in vitro studies^11–13^ and indicating an ongoing post-injury repair mechanism. In contrast, human IPF lungs depicted a divergent path, with AT2 cells branching into either AbBa cells or AT1 cells. The apparent disruption in the IPF lung regenerative process is in line with descriptions of AbBa cell persistence as an intermediate, non-regenerative state^5^ potentially driving the progressive, irreversible fibrosis in IPF, as opposed to the resolution of fibrosis following acute injury in the mouse model. These findings highlight key challenges in translating animal models to human disease and suggest that the acute BLM mouse model might offer valuable insight into alveolar regeneration. Application of SRT to the repeat BLM instillation model^61^, in which a more persistent, senescent Krt8+ transitional alveolar cell state has been identified^53^, could provide more insights into disease progression in the IPF lung.

Our study illustrates the potential for spatial transcriptomics to deepen our understanding of IPF pathology and offers rich datasets to further probe the complex cellular interplay in lung fibrosis. This work provides resolution of key mechanisms underpinning IPF and proposes a divergent cellular trajectory towards arrested regeneration in the human IPF lung, as a potential target for the discovery of novel disease modifying therapies.

## Methods

### Experimental methods

#### Human lung tissues and ethics declaration

IPF lung tissue was obtained from lung transplant patients. Human samples were acquired with approval by the local human research ethics committee (Gothenburg, Sweden; permit number 1026-15) and participants gave written informed consent prior to inclusion. Healthy lung tissue was obtained from deceased donors with no known lung disease, where samples were acquired with approval by the local human research ethics committee (Lund, Sweden; permit number Dnr 2016/317). All investigations were performed in accordance with the declaration of Helsinki.

All human tissues selected for analysis were collected from the peripheral lung. Fresh-frozen tissues were obtained from four HC subjects and from four IPF patients. For each IPF patient, three different tissues were collected representing areas of mild (“B1”), moderate (“B2") or severe (“B3”) fibrosis within the same donor, as determined by histological inspection of H&E-stained samples.

#### Mice and bleomycin challenge

Female C57BL/6NCrl mice (Charles River, Germany) were 8 weeks old on the day of arrival at AstraZeneca R&D Gothenburg (Sweden). After an acclimatization period of 5 days, mice were challenged with 30 μl bleomycin (Apollo Scientific, BI3543, Chemtronica Sweden; 40 µg/mouse) dissolved in saline or saline via oropharyngeal route administration. Lung samples were collected at day 7 or day 21 following bleomycin challenge. The timepoints were selected to encompass the early phase of inflammation and tissue remodeling (d7), and the subsequent phase of established tissue damage (d21). The mice were housed in Macrolon III cages with poplar chips (Rettenmeier & Söhne) as bedding material, shredded paper, gnaw sticks and a paper house. They were kept in a facility with 12 h/12 h light/dark cycle at 21 ± 1 °C, 55 ± 15 % relative humidity and had free access to food (R70, Lantmännen AB, Vadstena, Sweden) and tap water. Animal handling conformed to standards established by the Council of Europe ETS123 AppA, the Helsinki Convention for the Use and Care of Animals, Swedish legislation, and AstraZeneca global internal standards. All mouse experiments were approved by the Gothenburg Ethics Committee for Experimental Animals in Sweden and conformed to Directive 2010/63/EU. The present study was approved by the local Ethical committee in Gothenburg (EA000680-2017) and the approved site number is 31-5373/11.

#### Mouse tissue collection

Mice were anesthetized with isoflurane (5%, air flow ∼2 L/min), placed on the operating table, and maintained with 3% isoflurane (air flow ∼0.7 L/min). An incision was made in the skin from the middle of the stomach up to the chin. 0.1 mL heparin was injected through the diaphragm to the heart, and the abdomen aorta was cut to bleed the mice, followed by a cut in the apex of the heart. The heart and right lung lobes were tied off. The left lobes were collected and snap frozen for downstream analyses.

The pulmonary circulation was perfused via the pulmonary artery with 0.8 mL 37°C saline followed by 0.6 mL 37°C low-temperature melt agarose (SeaPlaque) solution. The lung was then inflated with 0.4-0.5 mL 37°C low melt agarose solution via the trachea and tied off. The lung was collected and snap frozen in pre-chilled NaCl over dry-ice, and stored at -80C for further analyses.

#### Generation of spatially resolved transcriptomics

OCT-embedded human lung tissue-blocks and agarose-inflated mouse lung tissues were cryosectioned at 10 µm (mouse) or 12 µm (human) thickness with the cryostat temperature set to -20°C and -10°C (mouse) or -15°C (human) for the specimen head. For the human lung samples, RNA quality was estimated through total RNA extraction from 10 tissue sections with a RNeasy Plus Mini kit (Qiagen). Thereafter RNA integrity number (RIN) was measured using a 2100 Bioanalyzer Instrument (Agilent) and ranged between 5.4 and >8, except for one sample (IPF donor 2, B3) with a RIN of 3. Despite lower RIN values for some tissues, after taking histological integrity into account they were chosen to be included for further analysis. For the mouse sections, 10 sections were stored in -80°C prior to RNA extraction using the Rneasy micro kit (Qiagen). RNA quality was assessed using a 5300 Fragment Analyzer (Agilent) and the RIN values were >9 for all mouse samples.

The lung tissue samples were cryosectioned onto the Visium Gene Expression slide. All slides were stored at -80°C until further processing. Tissue fixation and staining followed the Methanol Fixation, H&E Staining, and Imaging Visium protocol (10X Genomics). Stained human lung sections were imaged using the Axio Imager.Z2 (ZEISS) light microscope at 20X magnification, and thereafter stitched using Vslide (MetaSystems). Mouse lung sections were imaged at 20X magnification using an Aperio Digital Pathology Slide Scanner (Leica Biosystems).

Sequencing libraries were prepared according to the Visium Spatial Gene Expression User Guide (10X Genomics, Rev C). The human tissue sections were permeabilized for 15 min and amplification of cDNA was performed with 15-17 cycles and indexing with 12-14 cycles. The mouse lung sections were permeabilized for 15 min, and cDNA amplification and indexing were performed with 16-17 cycles and 8-15 cycles, respectively. Permeabilization times had been optimized prior to the experiments using the Visium Tissue Optimization kit.

The human sample libraries and the mouse sample libraries were pooled separately and sequenced. A 1% PhiX spike-in was included. The pooled libraries were loaded at 300pM onto a NovaSeq 6000 (Illumina) machine and sequenced on the S4 flowcell using the following set-up: Read1: 28 bp, Index 1: 10 bp, Index 2: 10 bp, Read2: 90 bp. A total of 255-444 M reads (avg. 349 M) and 151-571 M reads (avg. 325 M) per sample were generated for human and mouse, respectively.

#### Histopathology annotations

Histopathological assessments were performed on the Visium H&E-stained tissue sections using the Loupe Browser (10X Genomics) software. The data was manually annotated into major tissue compartments based on tissue morphology. The human lung data was classified into “blood vessel”, “large airway”, “diseased (remodeled) tissue”, “fibroblastic foci / fibrous tissue”, “inflammation”, and “within normal limits” (alveolar), where “inflammation” was distinguished as areas with large aggregations of immune cells and “diseased tissue” largely corresponded to clearly recognizable changes in normal lung architecture. The “fibroblastic foci / fibrous tissue” was distinguished based on their microscopic appearance, characterized by the density and shape of nuclei present, and increased amounts of collagenous matrix, consistent with the appearance of the fibroblastic foci found in IPF lungs. The mouse data was categorized into similar groups of “blood vessel”, “large airway”, “within normal limits” (alveolar), “inflammation (d7)”, “inflammation (d21)”, and “suspect fibrosis / fibroplasia (d21)”. The areas annotated as “inflammation (d7)” in lungs collected at day 7 were composed of both inflammatory and fibrotic tissue as they were indistinguishably mixed, while “inflammation (d21)” labelled dense immune cell aggregates. Thus, the spots labeled as both “inflammation (d7)” and “suspect fibrosis/fibroplasia (d21)” contains fibrotic tissue.

### Computational processing and analysis

#### Processing Visium sequencing data

Raw human and mouse sequencing data FastQ files were processed using the Space Ranger 1.2.2 (10x Genomics) pipeline. Sequencing reads were mapped to their respective reference genomes GRCh38 (human) and mm10 (mouse). H&E images were manually aligned to the fiducial frame and tissue-covered spots were identified using the Loupe Browser (v.6, 10X Genomics) software.

#### Mapping single cell types spatially with cell2location

In the human samples, spatial deconvolution was performed using cell2location^31^ against a previously published pulmonary fibrosis scRNA-seq dataset (GEO accession GSE135893)^5^ (referred to as the *Habermann (2020)* dataset). The cell2location method uses signatures from the provided scRNA-seq data to infer absolute numbers (density) of cell types within each spatial spot. The single-cell regression model was trained with max_epochs = 250 after selecting genes with parameters nonz_mean_cutoff = 1.25, cell_count_cutoff = 5, and cell_percent_cutoff = 0.05. The cell2location model was thereafter obtained with parameters max_epochs = 10000, detection_alpha = 20, and n = 7.

For the mouse data, a scRNA-seq dataset produced from the bleomycin-induced lung fibrosis mouse model collected at multiple time points (including d7 and d21) was used (GEO accession GSE141259)^11^ (referred to as the *Strunz (2020)* dataset). For spatial deconvolution, we used max_epochs = 400 for single-cell model generation using the parameters nonz_mean_cutoff = 1.10, cell_count_cutoff = 4, and cell_percent_cutoff = 0.02 for gene selection. For model training, max_epochs = 15000, detection_alpha = 20, and n = 7 was applied.

#### Downstream quality control and processing of Visium data

Data filtering, processing, and analyses of the Visium data were performed in R (v.4.0.5) using the STUtility (v.1.1.1)^62^ and Seurat (v.4.1.1)^63^ packages.

For the human IPF and HC samples, spots under the tissue were selected for downstream analysis, and the data was imported into R using the STUtility function ‘InputFromTable’ where initial gene and spot data filtering was performed by setting the minimum UMI count per spot to 350, minimum UMI count per gene to 100, minimum number of genes per spot to 10, and minimum number of spots per gene to 5. Spots were thereafter further filtered by content of mitochondria-associated genes, where spots with less than 30% was allowed, and content of blood contamination detected using hemoglobin gene expression, where spots < 30% were kept. Gene information was retrieved via biomaRt^64^ and used to select for “protein coding”, “IG” (immunoglobulin), and “TR” (T cell receptor) gene biotypes, as well as to flag genes positioned on the X and Y chromosomes for removal to avoid gender biases in the analyses. Post-quality control, an average of 4,043 spots per tissue section and across all sections was obtained, which yielded over 100,000 spots in total with data from over 15,000 genes for the human Visium dataset. Normalization and scaling of the data was performed using the ‘SCTransform’ function^65^ (Seurat package), specifying sample ID and donor as variables to regress out, to remove the major effects of technical and interindividual differences.

Visium data generated from mouse lungs was filtered in a similar manner, apart from omitting the number of genes per spot (“minGenesPerSpot”) cutoff when loading the data using ‘InputFromTable’, and an adjusted spot filtering for number of UMIs per spot set to 300. The final mouse Visium dataset included information of more than 15,000 genes in over 90,000 spots across all samples. ‘SCTransform’ was thereafter applied to the data, specifying the animal ID as a variable to regress out. All thresholds for filtering were set based on initial examination of the raw data to exclude low quality spots (or spots outside of tissue areas) and genes with low expression.

#### Differential expression analysis on Visium pseudo-bulk data

For initial differential gene expression analysis (DEA) between conditions, pseudo-bulk datasets were generated from the Visium gene count matrices. For the general condition comparison (HC vs IPF for human and vehicle control vs BLM d7 or d21 for mouse), this was achieved by aggregating the raw counts per gene across all spots belonging to a donor or animal. Thereafter, DESeq2^66^ was used for the differential gene testing by specifying “condition”, with “control” as reference, in the design. For the bulk comparison of fibrotic regions between IPF and BLM d7 or d21, pseudo-bulk data on a donor/animal level was obtained from the annotated tissue sections by pooling the counts from spots labelled as diseased (fibrotic, FF, remodeled, or inflamed (BML d7)) in the disease condition samples. Combined counts from the fibrotic regions were compared against the pseudo-bulk data from entire control samples using DESeq2 (with “condition” set as the design), for each species and/or timepoint separately. To compare results between species, orthogene^67^ was used to identify mouse gene orthologues of the human genes, and the DESeq results were filtered to include only genes with available orthologues and present in all datasets (total of 12611 genes).

#### Non-negative matrix factorization (NMF)

Deconvolution through NMF was applied to the Visium gene expression data using the ‘RunNMF’ function in STUtility. The factorization method decomposes the data into a set number of factors that are expressed as non-negative values (activity) within each data point (spot) along with a feature (gene) loading matrix, describing the contribution (weight) of each gene to the factors.

The full human (HC and IPF) dataset was deconvolved into 30 factors (“hsNMF”), while the mouse data (vehicle control and BLM) was split by timepoint (d7, d21) before each subset was deconvolved into 30 factors (mmNMF_d7_, mmNMF_d21_). To describe each factor, functional enrichment analysis of the top 25 most contributing genes for each factor was performed using the ‘gost’ function in the gprofiler2 (v. 0.2.1) R package^68^, with the “hsapiens” (human) or “mmusculus” (mouse) organism specified. All factors were further annotated by examining the top contributing genes, the spatial localization of factor activity, and their abundance in different samples (diseased or control).

To compare hsNMF and mmNMF_d21_ factors across species, the R package orthogene was first used for gene symbol conversion between human and mouse, and then the top 100 contributing genes for each factor was compared using Jaccard similarity index computation. Jaccard index was calculated as the intersection over the union of each gene set pair.

The distribution of each hsNMF factor within the human samples were estimated by counting the number of spots belonging to the 99^th^ percentile of factor-active (F^hi^) spots and computing their frequency versus the total number of spots in each biopsy category (B0-3).

Spatial co-localization of factors and cell types was estimated by computing the pairwise Pearson correlation coefficient between spot factor activity and inferred cell type density. To identify donor variability in co-localization, the human Visium data was split into groups of HC (all HC donors), IPF donor 1, IPF donor 2, IPF donor 3, and IPF donor 4, before computing the correlation scores.

The most active (99^th^ quantile) hsNMF F14 spots (denoted F14^hi^) were subclustered by conducting a principal component (PC) analysis and using PCs 1-8 as inputs for ‘FindNeighbors’ and ‘FindClusters’ (resolution = 0.4), which generated five clusters. The mmNMF_d21_ F14^hi^ spots were subclustered using the same approach, but with PCs 1-14 as input and clustering resolution set to 0.5, obtaining three clusters.

#### Radial distance analysis

Fluctuations in gene expression and cell type densities along a radial distance from the hsNMF F14^hi^C0 or mmNMF_d21_ F14^hi^C0 regions (region of interest; ROI) were computed. The distance information from each section containing the ROI was extracted using the *semla* R package^69^ (v. 1.1.6; R v. 4.2.3; Seurat v. 4.3.0.1) with the ‘RadialDistance’ function, where singletons were excluded in the human analysis. In the human IPF data, distance correlation coefficients were computed for the 1000 most variable genes at a 500 µm distance from the ROI border using Pearson correlation. P-values were corrected using the Benjamini-Hochberg (BH) method and used to filter for significant (adj. p < 0.01) genes. Cell type density correlation was obtained using Pearson correlation and BH-corrected p-values at a radial distance of 500 µm. Since a linear relationship may not be present in all cases, cell density and gene expression fluctuation as a function of radial distance was visualized using the ‘geom_smooth’ function (ggplot2) with method set to “gam” (generalized additive model) and formula “y ∼ s(x, bs = ‘cs’)”. For the mouse BLM d21 data, the cell type density across radial distance from the ROI was visualized using ‘geom_smooth’ with the “loess” (Locally Estimated Scatterplot Smoothing) method.

#### IPF fibrotic niche regulators and cell-cell communication

In the human IPF Visium data, the microenvironment surrounding hsNMF F14^hi^C0 spots was investigated by first identifying the nearest neighbors (using the ‘RegionNeighbours’ STUtility function) over two rounds, thereby including spots located ≤ 2 spot distances from hsNMF F14^hi^C0. Next, the selected neighboring (nb.) spots were clustered by first running PCA and then using PC 1-9 as input for ‘FindNeighbors’ and thereafter ‘FindClusters’ (resolution = 0.2), obtaining 6 clusters (nb. clusters 0-5). Marker genes were identified using ‘FindAllMarkers’ on the neighboring spot data subset and comparing each cluster against the remaining clusters. Due to their low abundancies, nb. clusters 3-5 were omitted in some of the downstream analyses.

Upstream regulators and active pathways for the nb. clusters (0-2) were predicted with Ingenuity Pathway Analysis (IPA; version 90348151, Ingenuity Systems, Qiagen), using the cluster marker gene lists (adj. p < 0.01). As a reference, marker genes for the hsNMF F14^hi^C0 cluster was also included in the analysis. These markers were generated by comparing hsNMF F14^hi^C0 spots against all other spots in the IPF Visium subset, using ‘FindMarkers’ with arguments “min.pct = 0.25” and “min.diff.pct = 0.1”. The output was thereafter compared using the R package multienrichjam (v. 0.0.72.900)^70^ and the top 20 upstream regulators and top 10 enriched pathways and diseases/functions were plotted.

Directional cell-cell communication analysis was employed within the spatially constrained nb. clusters using NicheNet (v. 1.1.1)^44^, a method in which ligand-target links are predicted using gene expression and a prior model that incorporates intracellular signaling. Information containing ligand-receptor interactions (“lr_network.rds”), ligand-target gene regulatory potential scores (“ligand_target_matrix.rds”), and weighted ligand-signaling and gene regulatory network (“weighted_networks.rds”) were retrieved from the NicheNet data repository (DOI: 10.5281/zenodo.3260758). Analyses were performed in four rounds based on which cluster(s) were specified as receiver and sender populations, 1) receiver: F14^hi^C0, senders: nb. clusters 0-5, 2) receiver: nb. cluster 0, sender: F14^hi^C0, 3) receiver: nb. cluster 1, sender: F14^hi^C0, and 4) receiver: nb. cluster 2, sender: F14^hi^C0. In all rounds, receiver genes were identified by setting the condition reference as data from all other spots not included in the analysis (of IPF and HC origin). The results from all four analyses were compiled and the top prioritized ligands (avg. correlation value > 0.075) sorted based on the round 1 results were used to visualize the corresponding results in the other analysis rounds and the gene expression levels across all selected clusters.

#### Spatial cell type compartmentalization in mouse

Cell type co-localization compartments were identified in the mouse Visium data using the Strunz (2020) cell2location results. Cell types annotated as “NA” and “low.quality.cells” were excluded and the Visium spot data was subset into groups of vehicle (d7 and d21), BLM d7, and BLM d21, before pair-wise correlations (Pearson) for each cell type across all spots within each subset were computed. Hierarchical clustering was performed, and compartments were defined based on a generally applied tree height (*h*) cut-off of 1.5. A Sankey diagram was drawn based on the cell types falling into each compartment for each data subset. To visualize the spatial localization of the BLM d21 compartments F, H, and G, spot-wise compartment scores were computed by summing the inferred cell type densities for all cell types belonging to each compartment.

#### Translational analyses of human and mouse aberrant basaloid clusters in a shared gene-space

Selected IPF and BLM d21 samples were chosen for the integrated analysis (IPF 3 B1-B3, IPF 4 B1-B3, BLM d21 animals 1-5), based on having more pronounced fibrosis and presence of identified AbBa cell-dense regions. Raw count data were filtered to include only genes with orthologous name conversions, identified by orthogene^67^. Subsequently, a new assay was created from this filtered data for separate normalization of human and mouse datasets. The two data sets were then integrated based on the shared genes using the anchor integration approach in Seurat (‘FindIntegrationAnchors’ followed by ‘IntegrateData’. Default parameters), with specified anchor features identified using ‘SelectIntegrationFeatures’. Marker genes for the hsNMF F14^hi^C0 or mmNMF_d21_ F14^hi^C0 clusters were thereafter identified separately with ‘FindMarkers’ and comparing against all other same-species spots, using the integrated genes.

The identified hsNMF F14^hi^C0 and mmNMF_d21_ F14^hi^C0 marker genes (Bonferroni adj. p < 0.05) were analyzed for upstream regulator and canonical pathway enrichment prediction in IPA. Results were compared across species using the R package multienrichjam (v.0.0.72.900)^70^ to pinpoint shared and unique upstream regulators and pathways. The most significant regulators (p value < 10^−7^, right-tailed Fisher’s exact test) and pathways (p value < 10^-4^) were visualized in clustered network (cnet) plots, which groups predicted molecules into clusters (“Nodes”), based on shared contributing marker genes.

#### Lymphocyte aggregate comparison

The human IPF Visium data was split based on donor, and processed separately by running SCTransform() and NMF, producing 30 new and more refined subject-specific factors for each IPF donor. Examining the spatial factor activity and gene contribution, it was possible to identify one factor for IPF donors 1-3 that corresponded to histological findings of lymphocyte aggerates. For IPF donor 4, no corresponding factor could be identified. The selected factors that exhibited a signature for dense lymphocyte accumulations were factors 10 (IPF donor 1), 15 (IPF donor 2), and 17 (IPF donor 3). For the mouse data, the NMF results produced for each time point was used, and factors 11 (day 7) and 12 (day 21) were identified as corresponding to tertiary structure-like (TLS-like) features. In the mouse day 21 NMF analysis, we moreover identified factor 13 adjacent to the activity of NMF_d21_ factor 12.

The top 100 most contributing genes for each of the identified factors were selected and their gene loadings were scaled between 0 and 1 (each factor separately). Only genes which were found to have orthologous gene names in both species (based on conversion using ‘orthogene’) were selected, and to reduce the set of genes for the visualization in Fig. 6b, genes with a summed scaled loading of > 0.5 were used.

Cell densities and detection rates were estimated in the spots with the highest (99^th^ percentile) factor activity. The inferred cell type densities produced using cell2location with the Habermann et al. (human)^5^ and Strunz et al. (mouse)^11^ datasets were used. All immune cell types were selected for evaluation, however, for mouse, the following cell types were excluded from the visualization as they did not exhibit a relevant signal and lacked comparable human cell types: “AM (BLM)”, “AM (PBS)”, “Non classical monocytes (Ly6c2-)”, “Fn1+ macrophages”, “M2 macrophages”, “Themis T cells”, and “T cell subset”. For each subject, the average cell density was measured as the average inferred cell density among the selected spots and the detection rate was calculated as the percentage of spots displaying a density score higher than 0.5.

#### Spatial cell co-localization trajectory analysis

Cell type densities, inferred using the Habermann (human IPF) or Strunz (mouse BLM) scRNA-seq datasets, for “AT2 cells”, “Activated AT2 cells”, “Krt8+ADI”, and “AT1 cells” (mouse) or “AT2”, “Transitional AT2”, “KRT5-/KRT17+”, “AT1” (human) were used. Spots with the highest abundancies (95^th^ percentile) of these cell types were selected and used as input for dimensionality reduction with UMAP (n.neighbors = 30, min.dist = 0.1, for both the mouse and human analyses). In parallel, the cell type densities were used to produce low resolution clusters using ‘FindNeighbors’ and ‘FindClusters’ (mouse: resolution = 0.2, human: resolution = 0.1), to identify a cluster that corresponded to the AT2-dense spots (AT2-cluster). Trajectory analyses using the Slingshot approach^71^ (v. 1.8) were then applied to each set of UMAP spot embeddings with the ‘getLineages’ function and assigning the AT2-clusters as starting points. Curves were extrapolated using ‘getCurves’ (approx_points = 300, thresh = 0.01, stretch = 0.8, allow.breaks = FALSE, shrink = 0.99), and thereafter visualized on top of the UMAP embeddings. Pseudo-time was estimated by passing filtered gene count data (genes detected (>5 transcripts) in at least 1% of the total number of spots) and Slingshot curves into a Negative Binomial Generalized Additive Model using the ‘fitGAM’ function from the tradeSeq R package (v. 1.4.0)^72^. In the human data, two curves were identified, and the visualized pseudo-time is the max value of the two pseudo-time curves.

## Acknowledgements

We sincerely thank the patients that donated lung samples, making this research possible. We further thank E. Sand for assistance with tissue sectioning and staining, G. Hamm for providing support on mouse lung tissue preparation, and S. Bates for running quality measurements on the mouse tissue. This work was financially supported by the Swedish Foundation for Strategic Research.

## Author contributions

J.H., M.S., A.O., G.B., S.J., P.L.S, and J.L., conceived the study; L.F., M.O.L., S.J, and M.S. planned and designed the experiments; T.V. and A.B. provided animals from the bleomycin mouse model; A.C., S.O., and M.O.L. collected the mouse lung tissue; S.J. and L.F identified and selected the human tissues; L.F. and M.O.L. carried out tissue sectioning and spatial gene expression experiments; J. Lindgren and M.O.L. sequenced the samples; L.S. performed histopathological annotations; B.K. processed the raw Visium data and performed cell type deconvolution; L.F. and M.O.L carried out computational analyses of the Visium data; Data interpretation by M.O.L., L.F., M.H., V.P., M.S., A.O., P.L.S., and J.H.; M.S., P.L.S., J.H., and A.O. supervised the project; L.F. created the final figures and illustrations; M.O.L. and L.F. drafted the manuscript with input from M.H., V.P., M.S., A.O., P.L.S., L.S., G.B., T.V., S.J., and J.H; All authors read and approved the manuscript.

